# Caf1 regulates Ash1 histone methyltransferase activity via sensing unmodified histone H3

**DOI:** 10.1101/2023.01.24.525315

**Authors:** Eojin Yoon, Ji-Joon Song

## Abstract

Histone modifications are one of key mechanisms to regulate gene expression. Ash1 is a histone H3K36 methyltransferase and involved in gene activation. Ash1 forms a large complex with Mrg15 and Caf1/p55/Nurf55/RbAp48 (AMC complex). Ash1 subunit alone has very low activity due to the auto-inhibition and the binding of Mrg15 releases the auto-inhibition. Caf1 is a scaffolding protein commonly found in several chromatin modifying complexes. Caf1 has an ability to sense unmodified histone H3K4 residue. However, the role of Caf1 in AMC complex has not been investigated. Here, we dissected the interaction among the AMC complex subunits, revealing that Caf1 uses the histone H4 binding pocket to interact with Ash1 near the histone binding module cluster. Furthermore, we show that H3K4 methylation inhibits AMC HMTase activity via Caf1 sensing unmodified histone H3K4 to regulate the activity in an inter-nucleosomal manner, suggesting that there is a crosstalk between H3K4 and H3K36 methylations. Our work reveals a delicate regulatory mechanism of AMC histone H3K36 methyltransferase complex.

## Introduction

DNA in eukaryotic cells forms into a high-order structure chromatin composed of a repeating unit of nucleosome. Chromatin structure is modified in various ways to precisely control gene expression and many protein complexes are involved in chromatin modifications. The N- or C-terminal tails of histones in nucleosome are targets of diverse covalent modification including methylation, acetylation, phosphorylation, ubiquitylation, and others (Strahl and Allis 2000). These modifications either directly control chromatin structure or serve as a platform to recruit other transcriptional regulators (Li et al. 2007). The covalent histone modifications are dynamically regulated by writers and erases, and recognized by readers via specific histone binding module such as PHD, Bromo, Chromo and WD40 repeat domains. Furthermore, there are crosstalks between the histone modifications. For example, ubiquitylation of histone H2A stimulates PRC2 histone H3K27 methyltransferase activity, and histone H3K4 methylation inhibits PRC2 activity.

Among these modifications, histone methylations play critical roles in activating or repressing gene expression. While H3K9 and H3K27 methylations are directly involved in gene repression, H3K4 and H3K36 methylations in gene activation. Ash1 in fly and ASH1L in human are histone methyltransferases (HMTase) and belong to Trithorax Group proteins. The histone target site of Ash1 had been controversial. However, it has been clearly demonstrated that Ash1 is a bona-fide histone H3K36 methyltransferase (An et al. 2011). The human homolog ASH1L has been linked to a number of diseases, such as Tourette’s syndrome, autism spectrum disorder, and intellectual disability (Zhu et al. 2016; Okamoto et al. 2017; Rongve et al. 2019; Shen et al. 2019; Satterstrom et al. 2020; Dhaliwal et al. 2021).

Ash1 is a multidomain protein containing AWS, SET, Post-SET, Bromo, PHD and BAH domains (Tripoulas et al. 1996; Callebaut et al. 1999). Except for the AT-hook domain, the majority of the conserved domains are concentrated in the C-terminal half of Ash1. AWS, SET and Post-SET domain forms a catalytic core and the Bromo, PHD and BAH domains are located in a proximity to form a histone binding module cluster (Fig 1A). Although the catalytic core is sufficient to methylate histone H3K36, it has very poor activity as the substrate binding pocket is blocked by an auto-inhibitory loop coming from the post-SET domain (An et al. 2011). Based on these findings, it was suggested that another element is required to release the auto-inhibitory loop of Ash1. Later studies found that Ash1 forms a stable complex with Mrg15 and Caf1 (a.k.a. Nurf55 and p55 for fly and RbAp48/46 in human), termed AMC complex (Huang et al. 2017; Schmähling et al. 2018). Mrg15 is a transcription factor containing Chromo and Mrg domains. Ash1 and Mrg15 complex shows full histone H3K36 methyltransferase activity. Consistent with this, the crystal structures of human ASH1L bound with MRG15 shows that the binding of MRG15 destabilizes the auto-inhibitory loop and in turn activates ASH1L histone methyltransferase activity (Hou et al. 2019; Lee et al. 2019). As Ash1 and Mrg15 complex without Caf1 is shown to have a similar HMTase activity to AMC full complex, the role of Caf1 in AMC complex remains unclear.

**Figure 1.**
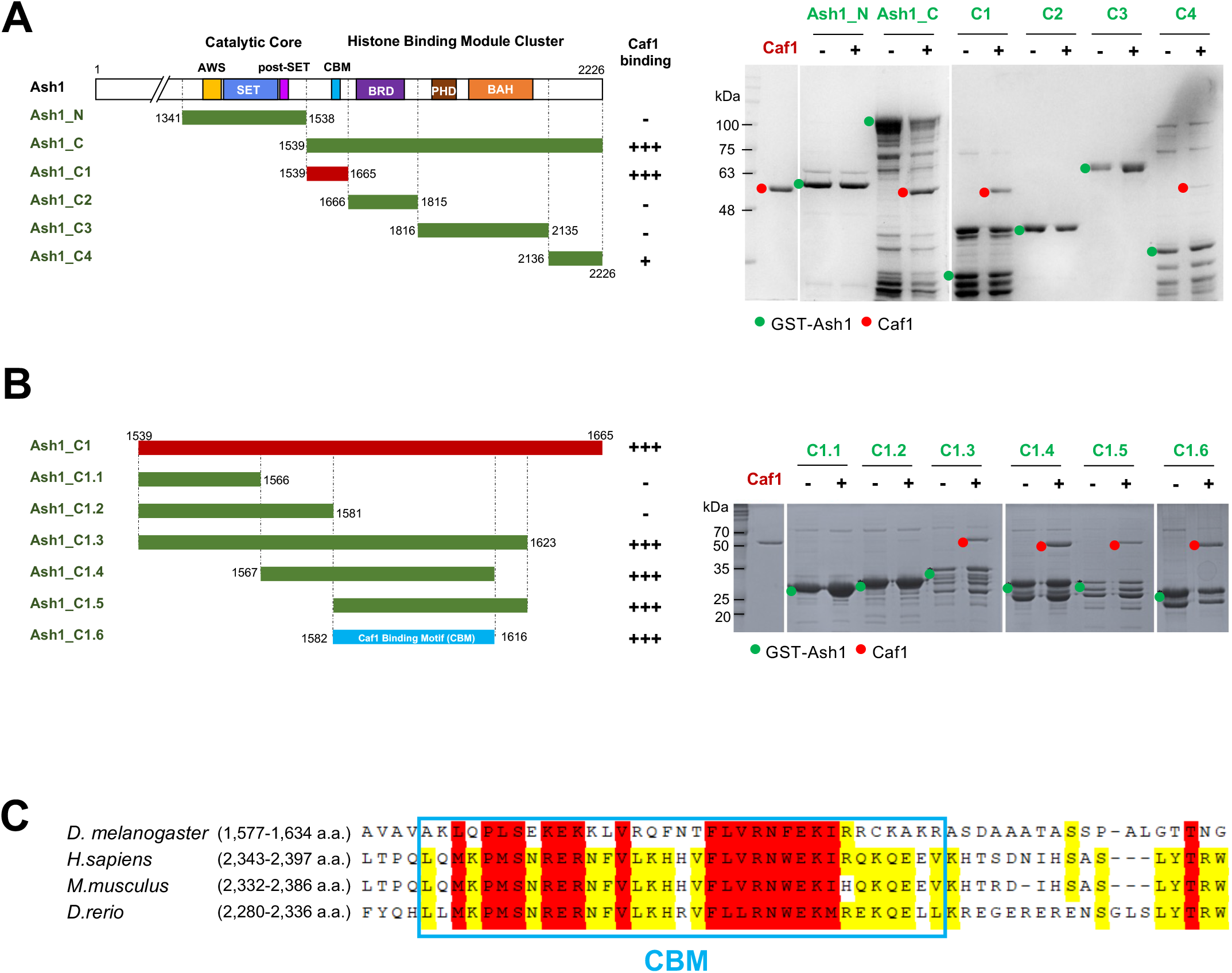
Caf1 binds to the proximal region of the Ash1 histone binding module cluster. **A**. A series of the truncation constructs of Ash1 and their binding to Caf1. Strong binding is indicated with ‘+++’, weak binding with ‘+’ and no binding with ‘-’ (left panel) based on GST-pulldown experiments with GST-Ash1 constructs and Caf1 (right panel). GST-Ash1 constructs are marked in green dots and Caf1 with red dots. **B**. Further truncation of Ash1_C1 and their binding to Caf1. The minimal region of Caf1 binding site on Ash1 is termed as ‘Caf1 Binding Motif (CBM)’. **C**. The sequence conservation of CBM in Ash1s among *Drosophila melanogaster, Homo sapiens, Mus musculus* and *Danio rerio*. Absolutely conserved residues are highlighted in red, and partially conserved residues in yellow.

Caf1 was initially identified as p55, a smallest subunit of Chromatin Assembly Factor 1 (CAF1) (Smith and Stillman 1989). Subsequently, Caf1 was identified as a key component of several chromatin modifiers including PRC2 histone methyltransferase, Mi-2/NuRD chromatin remodeler and Histone Acetyl Transferase complexes (Parthun et al. 1996; Taunton et al. 1996; Hassig et al. 1997; Zhang et al. 1997; Verreault et al. 1998; Czermin et al. 2002; Kuzmichev et al. 2002; Müller et al. 2002). Caf1 belongs to the WD40 protein family having a seven-bladed β-propeller structure. Caf1 has two conserved binding sites located on the top and the side of the β-propeller. Caf1 is shown to interact with histone H3, FOG-1 and Aebp2 using the top pocket, and with histone H4 helix 1, SUZ12, and MTA1 using the side pocket (Song et al. 2008; Lejon et al. 2011; Schmitges et al. 2011; Alqarni et al. 2014; Kasinath et al. 2018).

It is particularly interesting that Caf1 has preferential binding toward unmodified histone H3 tail and H3K4 methylation significantly lowered its binding to methylated histone H3 (Schmitges et al. 2011). Caf1 in PRC2 complex is shown to sense unmodified histone H3 tail to regulate PRC2 activity. Therefore, PRC2 shows significantly lower activity toward H3K4 methylated nucleosome.

However, the function of Caf1 in AMC histone H3K36 methyltransferase complex is not been investigated. Therefore, we explored the role of Caf1 in AMC complex. Specifically, we dissected the interaction between Caf1 and Ash1, showing that Caf1 utilizes the histone H4 binding pocket to interact with Ash1 near the histone reader domain cluster. Furthermore, we show that Caf1 in AMC also plays a role in sensing unmodified histone H3 to regulate AMC activity in an inter-nucleosomal manner. Our data revealed a delicate regulatory mechanism of AMC histone H3K36 methyltransferase complex.

## Results

### Caf1 interacts with the proximal region of the histone binding module cluster in Ash1

Ash1 directly binds to Mrg15 via the N-terminal region of Ash1 catalytic domain and it also binds to Caf1 in AMC complex (Hou et al. 2019; Lee et al. 2019). However, it has not been investigated how Caf1 binds to Ash1. Therefore, we dissected the interaction between Caf1 and Ash1. To identify the Caf1 interaction region on Ash1, we generated a series of truncated constructs of Ash1 and examined their interactions with Caf1 (Fig 1). First, we divided Ash1 into the N-terminal part (Ash1_N) containing the catalytic domain and the C-terminal part containing the histone binding module cluster (Ash1_C). GST-pulldown experiments show that Caf1 binds to Ash1_C. This data reveal that Ash1 uses two distinct regions for interacting Mrg15 and Caf1 (Fig 1A). To further dissect the Caf1 interaction site in Ash1, we generated four constructs and performed GST-pulldown assay (Fig 1A). They are a N-terminal region to the histone binding module (Ash1_C1), a bromodomain (Ash1_C2), a PHD-BAH domain (Ash1_C3) and a C-terminal region (Ash1_C4) to the histone binding module. We found that Caf1 strongly binds to the N-terminal region to the histone binding module cluster (Ash1_C1) and weakly associates with the C-terminus of the reader domains (Ash1_C4). To narrow down the interaction site in amino acid level, we then continued to trim down the Ash_C1 fragment into smaller fragments (Fig 1B). Among these fragments, Caf1 binds to 1,582-1,616 fragment (Ash1_C1.6). In addition, further trimming down the Ash1_C1.6 fragment substantially abrogate the interaction with Caf1 (Fig S1). Therefore, we concluded that Caf1 interacts with the region of 1,582-1,616 a.a. in Ash1 and we named this minimal binding region for Caf1 as ‘Caf1-binding-motif (CBM)’. We examined the sequence conservation of CBM among species (Fig 1C). The CBM region is highly conserved among human, mouse and zebrafish and fly, suggesting that Caf1 interact with Ash1 via CBM in other species as well (Fig 2S).

**Figure 2.**
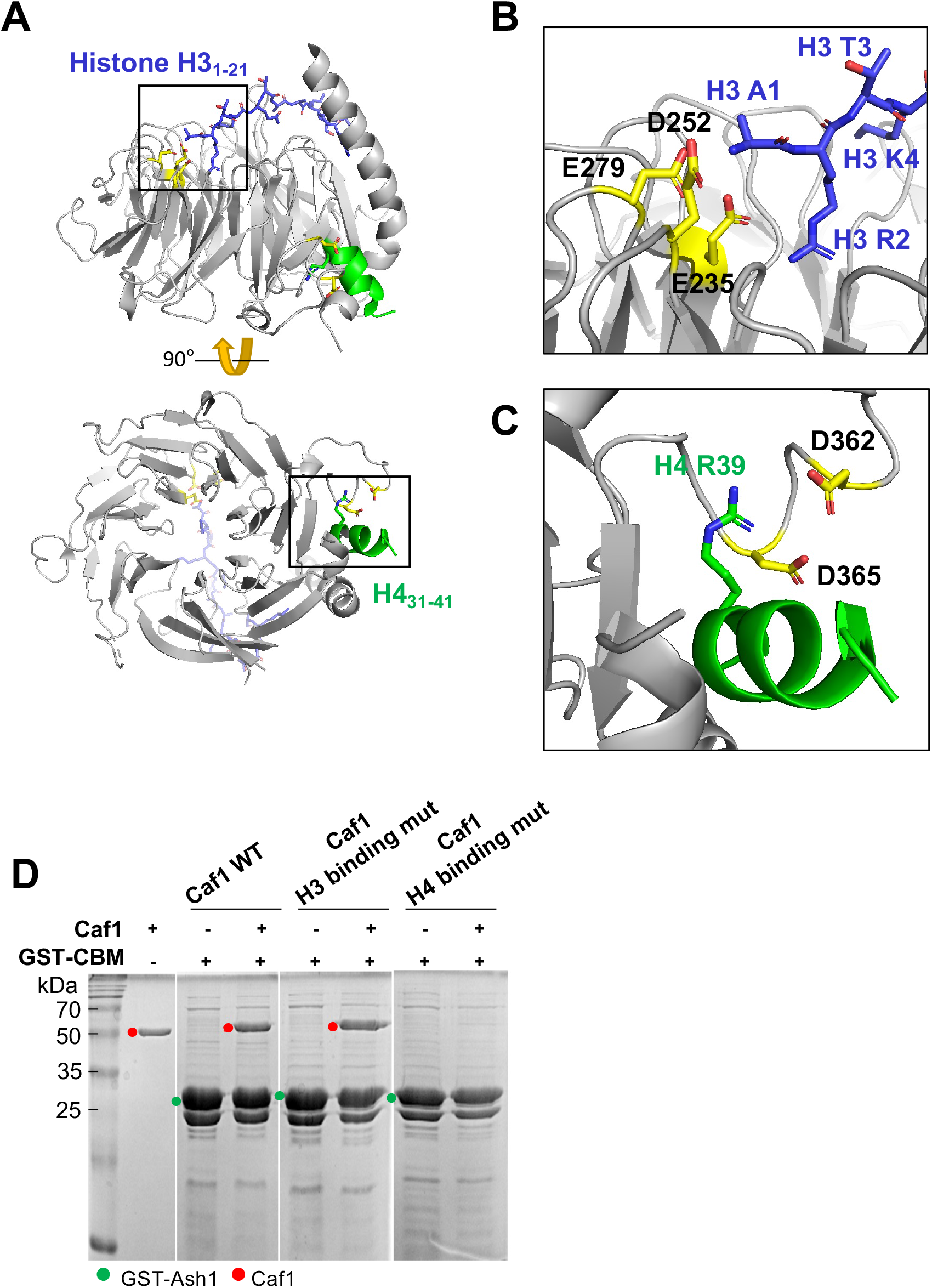
Caf1 utilizes the H4 binding pocket to interact with Ash1_CBM. **A**. The crystal structures of Caf1-histone H3_1–21_ complex (blue, PDB ID: 2yba and Caf1-H4 helix3_1-41_ (green, PDB ID: 3c9c) is shown in one Caf1. **B**. The histone H3 binding pocket is located at the central pore of Caf1. A1 and R2 residues of H3 N-terminal tail are coordinated with E235, D252 and E279 of Caf1. **C**. Caf1 utilizes a pocket located at the side of the WD40 β-barrel structure for H4 binding. The first helix of histone H4_31-41_ is accommodated with the carboxyl groups of D362 and D365, and the backbone carbonyl groups in a loop (366-371 a.a.) in Caf1. **D**. GST pulldown assay using GST-CBM and Caf1 and Caf1 mutants. Caf1 H3 binding mut has mutations of E235Q/D252K/E279Q in Caf1, and Caf1 H4 binding mutant jas mutations of D362A/D365A in Caf1.

### Caf1 utilizes the histone H4 binding pocket for Ash1 interaction

Caf1 has two histone binding pockets. One recognizing histone H4 is located at the side of WD40 β-propeller structure and the other recognizing histone H3 tail located at the central pocket (Fig 2A). The histone H4 binding pocket is shown to be very versatile as this pocket also binds to several other proteins including SUZ12 and MTA1 besides histone H4 (Schmitges et al. 2011; Alqarni et al. 2014). To examine whether Caf1 utilizes these pockets for Ash1 binding, we generated Caf1s where the histone binding pockets were mutated. First, we mutated histone H3 binding surface (E325Q/D252K/E279Q) (Fig 2B) and examined its binding to the Ash1 CBM fragment (Fig 2D). The mutation in the histone H3 binding pocket of Caf1 did not abrogate the interaction with Ash1. Next, we generated Caf1 mutant where the histone H4 binding pocket was mutated (D362/D365) (Fig 2C). This Caf1 mutant completely lost the binding to Ash1 (Fig 2D). These data show that Caf1 utilizes the histone H4 binding pocket to interact with Ash1.

To further examine the binding mode of Caf1 and CBM, we decided to analyze the Caf1 and CBM interaction using AlphaFold structure prediction. We generated a model of Caf1-CBM complex by feeding Caf1 and CBM sequence to AlphaFold structure prediction platform (Mirdita et al. 2022). Interestingly, the AlphaFold predicted structure shows that CBM forms a helix and binds to the histone H4 binding pocket of Caf1 (Fig 3A). In addition, the predicted structure of human ASH1L CBM and RbAP48 shows a similar binding mode to fly CBM-Caf1 interaction (Fig S3). These modeled structures are consistent with our GST-pulldown assay showing that the mutation of this histone H4 binding pocket abrogates the Caf1 and Ash1 interaction (Fig 2D). In the Caf1-histone H4 complex structure (Song et al. 2008), R39 of histone H4 is coordinated by two aspartates (D362 and D365) and the backbone carbonyl groups in the loop (366-371 a.a.). Interestingly, the highly conserved R1604 of Ash1 is coordinated with the same residues including D362 and D365 (Fig 3B). In addition, other absolutely conserved residues (E1591, V1595, F1601, L1602, N1605 and K1608) among species seems to be involved in the interaction with Caf1 (Figs 1C and Fig S4). To confirm this interaction mode predicted by the AlphaFold, we generated CBM_E1591A_ or CBM_R1604_ mutants and examined their binding to Caf1. GST-pulldown assay shows that these mutants do not bind to Caf1 (Fig 3A). These data shows that Caf1 utilizes the versatile histone H4 binding pocket for interacting with Ash1.

**Figure 3.**
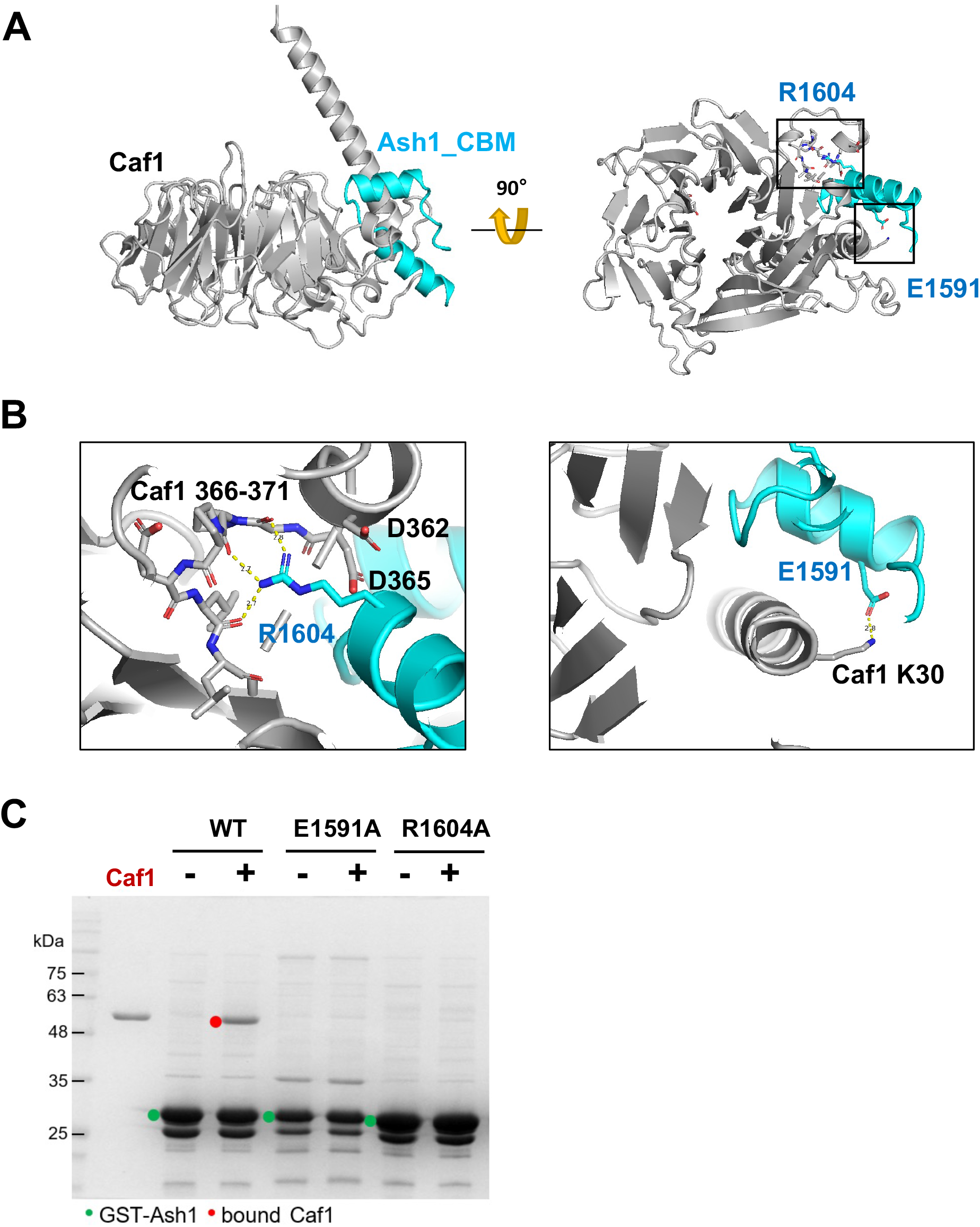
The binding mode CBM on Caf1. **A**. An AlphaFold predicted structural model of Caf1 (grey) bound to CBM (1,582-1,616 a.a., cyan). The CBM binds to Caf1 histone H4 binding pocket, forming a clip-like structure. **B**. Detailed interactions between Caf1 and CBM. R1604 of CBM makes extensive electrostatic interaction with carboxyl groups in the loop (366-371 a.a.) and D362/D365 residues in Caf1 (left panel). In addition, E1591 in CBM makes a salt bridge with K30 in Caf1, further stabilizing the Caf1_CBM interaction. **C**. GST pulldown assay using GST-CBM wild-type and mutants for Caf1 binding. GST-CBMs are marked with green dots, and CBM bound Caf1 in red dots.

### Caf1 senses unmodified histone H3 tail to regulate AMC HMTase activity

Caf1 subunit (RbAp48) of PRC2 functions as a sensor for active chromatin marks by recognizing unmodified histone H3. Caf1 has preferentially binding toward to unmodified histone H3 tails and methylated histone H3 (H3K4_me3_) inhibits PRC2 mediated histone H3K27 methylation (Schmitges et al. 2011). Therefore, we sought a possibility that Caf1 subunit in AMC functions in a similar manner to PRC2. To test this hypothesis, we measured the HMTase activity of AMC with unmodified or H3K4 methylated G5E4 nucleosome arrays. For H3K4 methylated nucleosome, we installed methylated lysine using the methyl-lysine analogue (MLA) protocol (Fig S5) (Simon et al. 2007).

Interestingly, AMC shows significantly lower activity for methylated nucleosomes than with unmodified nucleosomes, suggesting that Caf1 in AMC senses unmodified histone H3 as it does in PRC2 (Fig 4A). As Caf1 interacts with unmodified histone H3 tail utilizing the central pore of the WD40 β-barrel structure, we examined whether the H3 binding pocket of Caf1 is utilized to sense unmodified histones in AMC. We generated mutations (E235Q_D252K_E279Q) in the histone H3 binding surface of Caf1 (Fig 2B) to abolish the interactions between Caf1 and histone H3. We then measured the AMC HMTase activity. Contrast to wild-type AMC, AMC having the Caf1 H3 binding mutant shows similar activity for both unmodified and methylated nucleosome array (Fig 4B). Furthermore, we assessed the quality of the two nucleosome arrays to confirm that this observation is not due to the different quality of the nucleosome arrays using DOT1L histone methyltransferase. DOT1L activity was not affected by the presence of methylation in histone H3K4 (Fig 4C). These data imply that Caf1 subunit in AMC senses unmodified histone H3K4.

**Figure 4.**
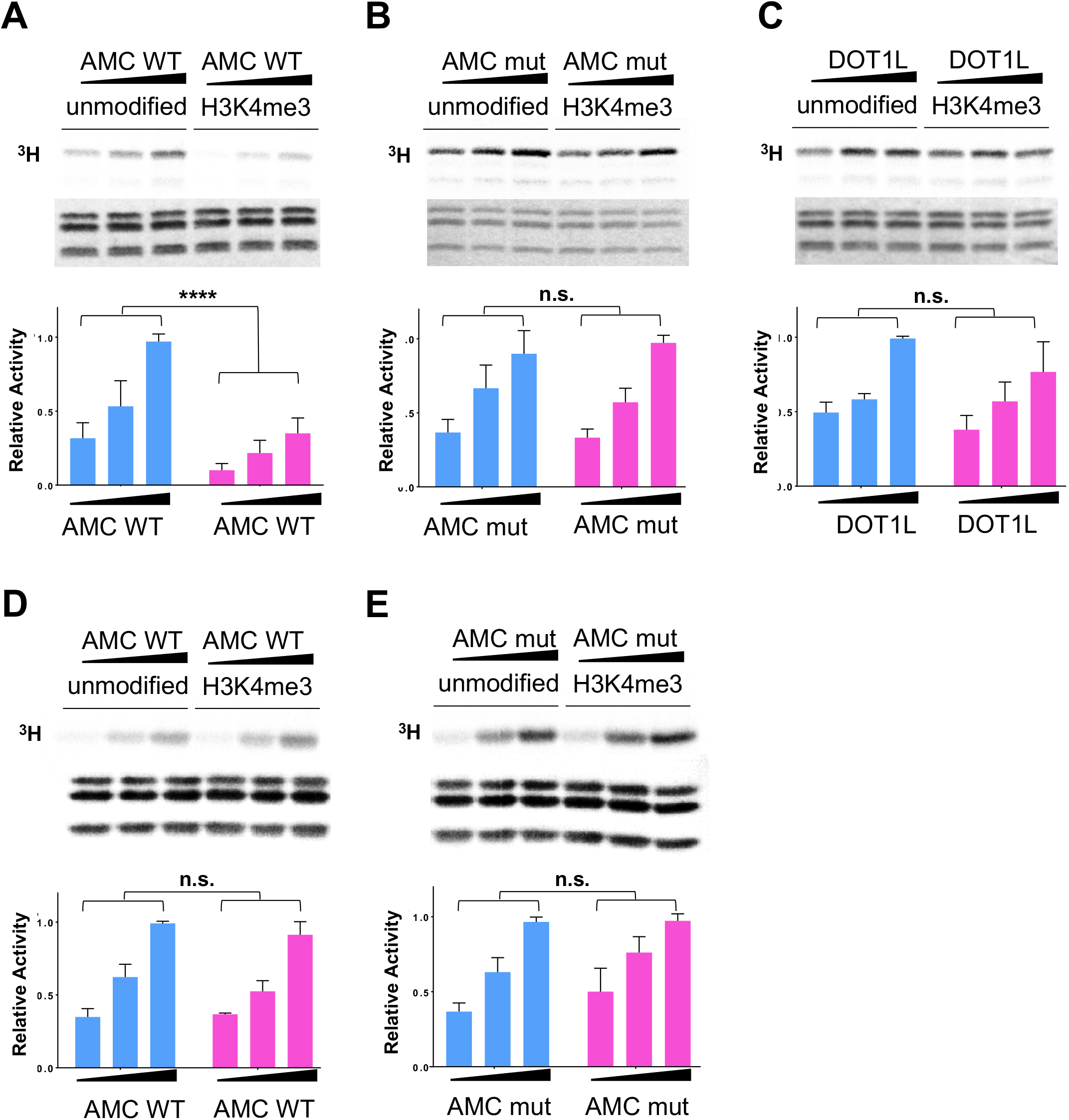
Caf1 in AMC complex senses unmodified histone H3 on nucleosome array. **A.-C**. HMT assay using G5E4 nucleosome array and AMC complex (70, 100 and 140 nM) (top panels). The relative activity of AMC complexes on unmodified nucleosome array is shown in blue, and H3K4me3 nucleosome array in magenta (bottom panels): AMC wild-type (AMC WT), AMC with Caf1 H3 binding mutant (E235Q/D252K/E279Q) (AMC mut) and DOT1L (n=3, **** – p value < 0.0001, n.s.=not significant). **D., E**. HMT assay using mononucleosome with 80 bp linker DNA (top panels) and the relative activity shown in a bar graph (n=3, n.s.=not significant) (bottom panels).

Recently, PRC2 is shown to work on dinucleosomes (Poepsel et al. 2018). Therefore, we examined whether AMC works on dinucleosomes and the inhibitory effect of methylated histones on AMC results from inter-nucleosomal phenomena. We generated unmodified and H3K4 methylated mononucleosome with 147 bp Widom 601 DNA with 80 bp linker DNA at one site and measured the AMC HMTase activity. Both wild-type AMC and the histone H3 binding mutant AMC show no significant difference for unmodified and methylated mononucleosomes (Figs 3D and 3E). The fact that AMC shows similar activity to both unmodified and methylated mononucleosomes suggests that AMC acts on more than one nucleosome and the inhibitory effect of methylated nucleosome on AMC activity results from inter-nucleosomal interactions.

## Discussion

In this study, we dissected the interaction among the subunits of fly Ash1 histone methyltransferase complex, showing that Caf1 subunit interacts with the highly conserved region named ‘Caf1-Binding-Motif (CBM)’ at the N-terminal proximity to histone binding module of Ash1 via the Caf1 histone H4 binding pocket. Furthermore, we demonstrated that Caf1 in Ash1 complex plays a role in sensing unmodified histone H3 to regulate the histone methyltransferase activity through inter-nucleosomal interaction.

Ash1 has a histone binding module cluster composed of bromodomain (BRD), PHD and BAH. Interestingly, our data showed that Caf1 binds to CBM located at N-terminal proximity to the histone binding module cluster (Figure 1). Considering that Caf1 is also a histone binding module, Caf1 together with the BRD-PHD-BAH cluster may function cooperatively to recognize combinatorial histone modifications. CBM is highly conserved among species, suggesting that Ash1s from other species also utilizes CBM motif for Caf1 interaction (Figs 1C).

Caf1/p55/Nurf55/RbAp48 is a very versatile protein, which functions as a scaffold to assemble large chromatin modifying complexes and as a histone binder. Our biochemical and structural analysis shows that Caf1 utilizes the histone H4 binding pocket located at the side of β-propeller structure to interact with the CBM of Ash1. As this pocket is also shown to interact with several other proteins such as SUZ12 and MTA1 (Fig S3), it is likely that there are more other proteins interacting with this pocket.

Caf1 also binds to histone H3 tails using the pocket at the top of β-propeller structure. At the histone H3 binding pocket, the *ε*-amine group of H3K4 is coordinated by two aspartates (E130 and E183). Caf1 has a nanomolar binding affinity (~ 600 nM) toward unmodified histone H3 and a hundred-fold lower affinity (>70 uM for H3Kme3) toward methylated histone H3 tails (Schmitges et al. 2011). This preferential binding ability of Caf1 as a component of PRC2 plays a critical role in sensing unmodified nucleosome substrate. Therefore, PRC2 has substantially lower activity toward H3K4 methylated nucleosome substrate. Interestingly, our data show that AMC histone H3K36 methyltransferase also shows substantially lower activity toward H3K4 methylated nucleosome than unmodified nucleosome, and that AMC complex containing Caf1 with the histone H3 binding pocket mutant does not discriminate methylated and unmethylated nucleosomes, implying that Caf1 in AMC complex functions a similar manner to PRC2.

Both histone H3K36 and H3K4 methylation marks are largely considered as active marks for gene expression. Therefore, it is counterintuitive that histone H3K4 methylation inhibits AMC H3K36 methylation activity. However, some of H3K4 methylation pattern in genome is quite distinct and does not overlap with H3K36 methylation pattern (Dorafshan et al. 2019). Therefore, it is plausible that Caf1 in AMC may play a role in discriminate these regions by sensing H3K4me3 free nucleosome.

Ash1 has a histone binding module cluster containing BRD, PHD and BAH domains, and Caf1. It is likely that these cluster recognizes combinatorial histone modifications. However, the full spectrum of the binding specificity of these domains are not clear. A large-scale structural analysis suggests that ASH1L Bromo domain interacts with multiple acetylated histones including H3K56ac and H4K59ac (Filippakopoulos et al. 2012). However, the biological role of the interactions has not been investigated. In addition, a recent structural analysis on ASH1L PHD domain shows that it binds H3K4 methylated peptide with very week affinity (100-200 uM range) (Yu et al. 2022) and does not bind to unmodified H3K4 peptide. It is quite interesting that Caf1 has strong affinity toward unmodified H3K4 and very week affinity to methylated H3K4, while PHD has very week affinity to methylated H3K4, and Caf1 and PHD are located in proximity. To add more complexity, the chromodomain of Mrg15, which is a component of AMC, is shown to bind to methylated histone H3K36 (Zhang et al. 2006). Structural study combined with functional assay will be required to precisely understand the contribution of histone modifications to Ash1 activity.

Chromatin modifying complexes often works on more than one nucleosome. For example, PRC2 complex is engaged in two nucleosomes (Poepsel et al. 2018). One nucleosome serves as a substrate nucleosome and the other nucleosome as an activating nucleosome whose modification stimulates PRC2 activity. Our data shows that the substrate preference of AMC for unmodified nucleosome was only observed when oligonucleosomes were used but not mononucleosome (Figure 4D). Therefore, the inhibitory effect on AMC activity by the methylated nucleosome likely occurs in an inter-nucleosomal manner. The domain organization in Ash1 is also consistent with the idea that Ash1 engages two nucleosomes. A large flexible region is linking the catalytic domain and the histone binding module cluster. It is imaginable that the catalytic domain binds to one nucleosome as a substrate and the cluster sense the other nucleosome, which plays a regulatory role.

The activity of Ash1 HMTase seems to be delicately regulated by several factors. First, the catalytic activity is regulated by the auto-inhibitory loop. Second, AMC complex contains a histone binding module cluster, which may recognize specific modifications to regulate the AMC activity. Thirds, this study reveals that Caf1 subunit senses unmodified histone H3 tail to regulate the activity.

Overall, our data imply that the activity of AMC histone H3K36 methyltransferase is highly regulated and further in-vitro and in-vivo studies are required to fully understand how AMC activity is regulated.

## Materials and Methods

### Protein expression and purification

Fly Caf1 wild-type, H3 binding pocket mutant (E235Q/D252K/E279Q), and H4 binding pocket mutant (D362A/D365A) mutant were cloned into a modified pFastBac_HTB vector having a 6x-His tag and TEV cleavage site on its N-terminus. It was expressed, purified from Sf9 cells using baculovirus system. After transfection, cells were harvested in 300mM NaCl, 50mM Tris-HCl (8.0), 5% glycerol buffer and lysed by 4 times of freeze-thawing. Lysed cells were cleared by 15,000rpm centrifugation for 2 hours. Equilibrated Ni-NTA resin (Qiagen) in same buffer with cells were subjected to cleared lysates for 2 hours, then washed with 5 cv of 1M NaCl and 2cv of 300mM NaCl, with 50mM Tris-HCl(8.0), 20mM imidazole buffer. Resin-bound Caf1 were eluted by 100mM imidazole elution buffer. 6x His-tag on Caf1 was cut by TEV protease, then Caf1 purification followed by HiTrap Q HP (Cytiva) and Superdex 200 26 600 (Cytiva) in the buffer containing 150mM NaCl and 50mM Tris-HCl (8.0). For the purification of AMC complex, 6x His-tagged Ash1 1041-2226, Mrg15 full length, and Caf1 full length were co-infected in Sf9 cells using baculovirus, and the cell was harvested in 300mM NaCl 50mM Tris-HCl (8.0) 5% Glycerol buffer. Cells were lysed by 4 times of freeze-thaw cycle, and cleared by 15,000rpm centrifugation for 2 hours. Equilibrated Ni-NTA resin (Qiagen) were incubated in cleared lysate for 2 hours and washed with 300mM – 1M (3cv) – 300mM NaCl buffer with 50mM Tris-HCl (8.0) 20mM imidazole buffer. Bound protein were eluted with 300mM NaCl, 50mM Tris-HCl (8.0) 5% glycerol 100mM imidazole buffer, and the affinity tag were cut by TEV protease. Cut His-tag and TEV protease were removed by recapture with Ni-NTA bead. Purification of AMC protein then followed by HiTrap Q HP (Cytiva) and Superdex 200 26 600 (Cytiva) in the buffer containing 300mM NaCl and 50mM Tris-HCl (8.0)

### Nucleosome reconstitution

Recombinant *Xenopus laevis* histone and 601 positioning DNA sequence with 80bp extension on one side (+80 DNA) were expressed and purified from Escherichia coli BL21 (DE3) pLysS and DH5a respectively, and remaining purification procedures were followed by published protocol (Luger et al. 1999). For the reconstitution of histone octamer, four types of histones were dissolved in unfolding buffer and mixed with H2A : H2B : H3 : H4 = 1.1 : 1.1 : 1 : 1 ratio in molar concentration. Buffer was changed to 2M NaCl solution through dialysis overnight at 4°C, and histone H2A/H2B dimer was discarded through Superdex 200 size exclusion column (Cytiva). After reconstitution of histone octamer, purified 601 DNA and octamer were mixed in between 1.0 to 1.6 ratio depending on the types of DNA or octamer. Salt concentration in mixture dropped slowly from 2M NaCl to 250mM NaCl by using dual peristaltic pump, and finally 0mM NaCl by dialysis.

### GST-pulldown assay

For GST pulldown assay, Ash1 fragments were cloned in pGEX-4T-1 vector. Cloned GST-tagged Ash1 fragments were expressed in *E.coli* BL21 (DE3) RILP strain at 18°C in the presence of 0.5mM IPTG for 18 hours, and harvested in 150mM NaCl 50mM Tris-HCl (8.0) buffer. Cells were lysed by sonication and cleared by centrifugation in 13,300 rpm for 10min. Equilibrated GST resin (Qiagen) was introduced to cleared lysate and incubated for 2 hours to bind with GST-bound Ash1 fragments, then washed several times with 150mM NaCl 50mM Tris-HCl (8.0) buffer. After washing, 0.1mg/ml of purified Caf1 200ul was added to GST resin and incubated for 1hour, and the unbound were washed out. Resin bound proteins were analyzed by SDS-PAGE.

### Methyltransferase assay (HMTase assay)

For HMTase assay, 1uM of wild-type AMC and mutant AMC were mixed with 100mM NaCl, 50mM Tris-HCl (8.0) and 150ng/ul of G5E4 nucleosome array or 1uM of +80 mononucleosome. Tritium SAM (PerkinElmer) were primarily mixed with 10x HMTase buffer (500mM Tris-HCl (9.0), 50mM MgCl2, 40mM DTT) and added to reaction mixture. Reaction was done in 25°C for 40minutes, stop by heating to 65°C for 5minutes, then SDS buffer were added and boiled in 95°C for 5minutes. Proteins were separated by 16% SDS PAGE and transferred to PVDF membrane by semi-dry transfer. Membrane was exposed to imaging plate for overnight, and the tritium signals were detected by FUJI-BAS. Intensities of detected band were measured by ImageLab program.

## Acknowledgements

We thank the member of Song Lab for helpful discussion and Yumi Shin for technical supports. This work was partially supported by grants (NRF-2020R1A2B5B03001517 and NRF-2019K1A3A1A18116012 to J.S.) by Korea National Research Foundation.

## Author contributions

EY and JS conceptualized the idea, EY performed the experiments, JS supervised experiments and EY and JS examined the data and wrote a paper.

## Conflict of Interest Statement

J.S. is a co-founder and CTO of Epinogen Co. Ltd.

---

**Figure S1.**
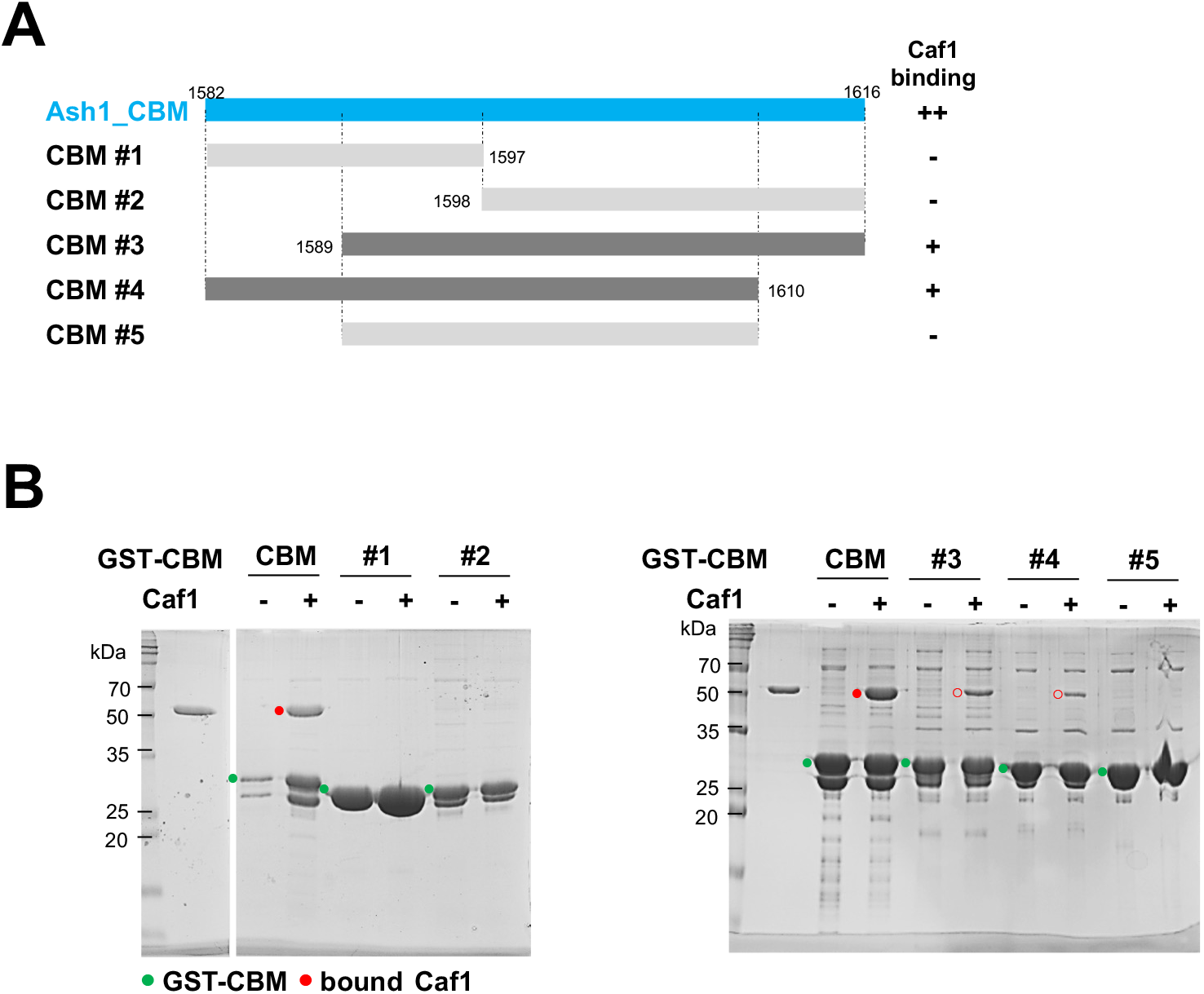
Caf1 Binding Motif (CBM) is a minimal binding site of Caf1. **A**. A series of the truncation constructs of CBM. Strong binding is indicated with ‘++’, weak binding with ‘+’ and no binding with ‘-’ based on GST-pulldown experiments. **B**. GST pulldown assay using GST-CBM fragments and Caf1. GST-tagged CBMs are marked in green and Caf1 marked with red dots. Weakly binding Caf1s are marked with red empty dots.

**Figure S2.**
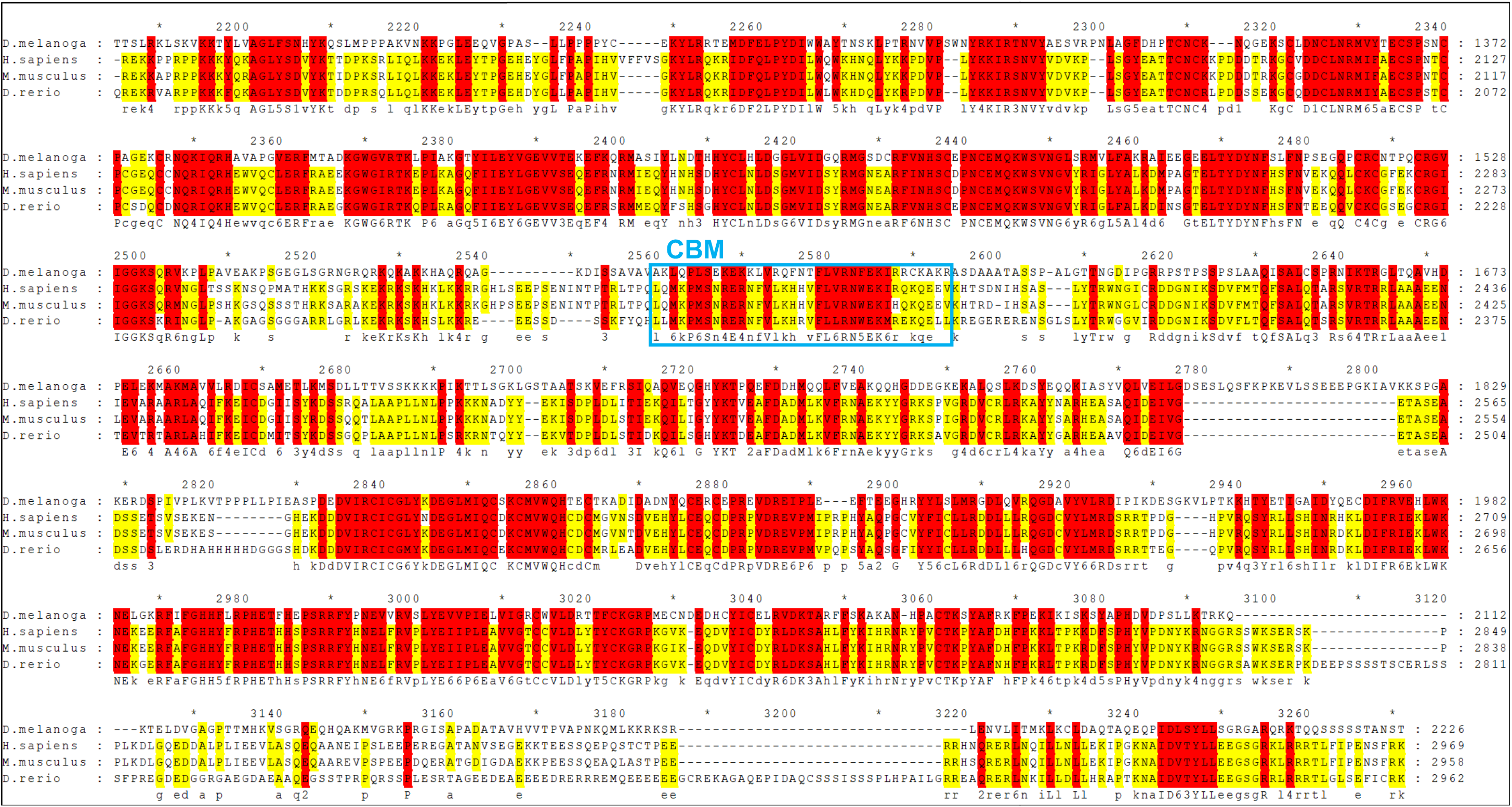
Sequence alignment of Ash1. The fly Ash1 1,227-2,226 sequence conservation among *Drosophila melanogaster, Homo sapiens, Mus musculus* and *Danio rerio*. Absolutely conserved residues are highlighted in red and partially conserved residues in yellow.

**Figure S3.**
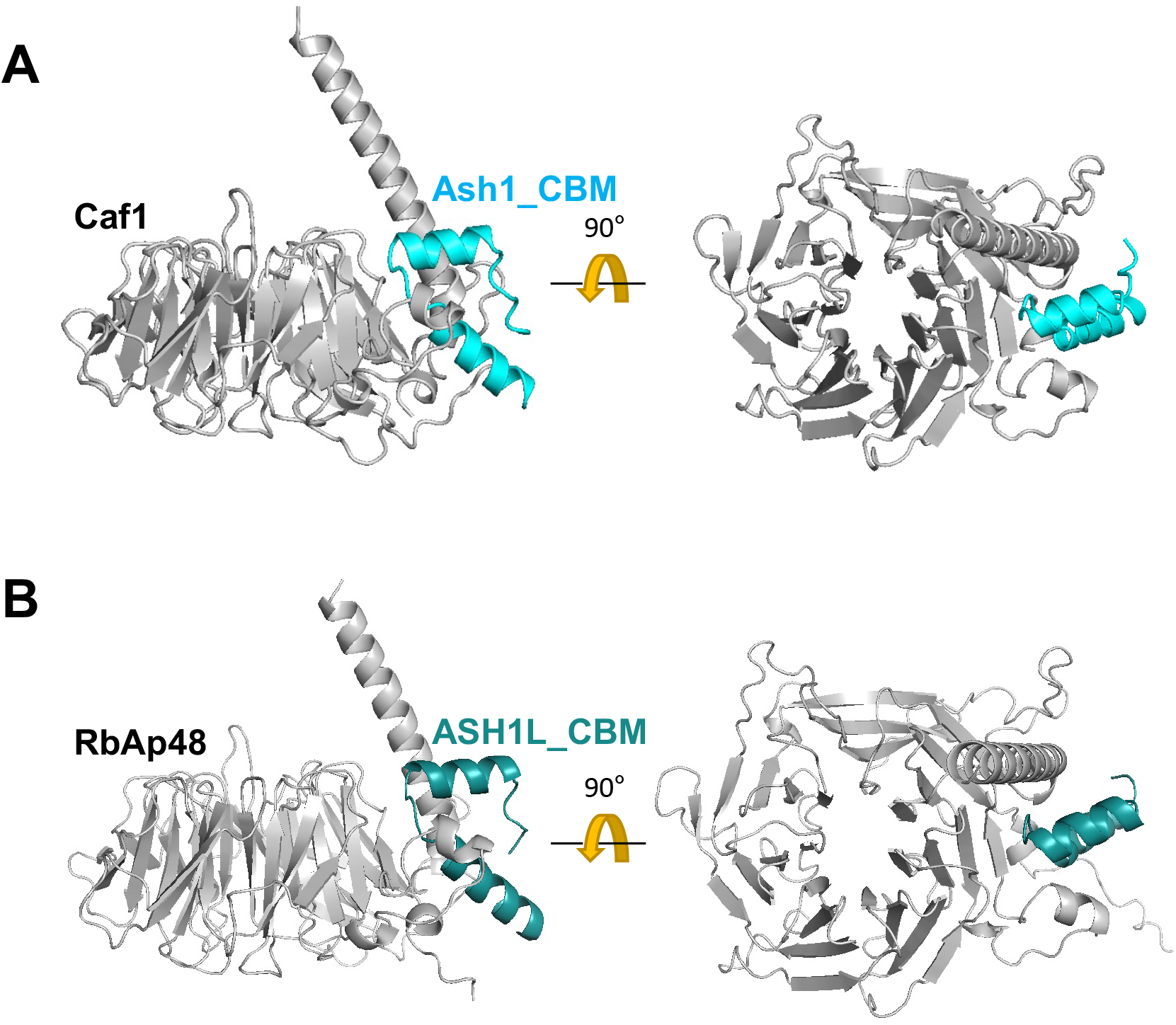
Alphafold predicted model of Caf1 and Ash1_CBM. **A**. Alphafold predicted model of Drosophila melanogaster Caf1 and Ash1_CBM sequence. Caf1 colored in grey, and Ash1_CBM colored in cyan. **B**. Alphafold predicted model of Homo sapiens RbAp48 and ASH1L_CBM sequence. RbAp48 colored in grey, and ASH1L_CBM colored in dark cyan.

**Figure S4.**
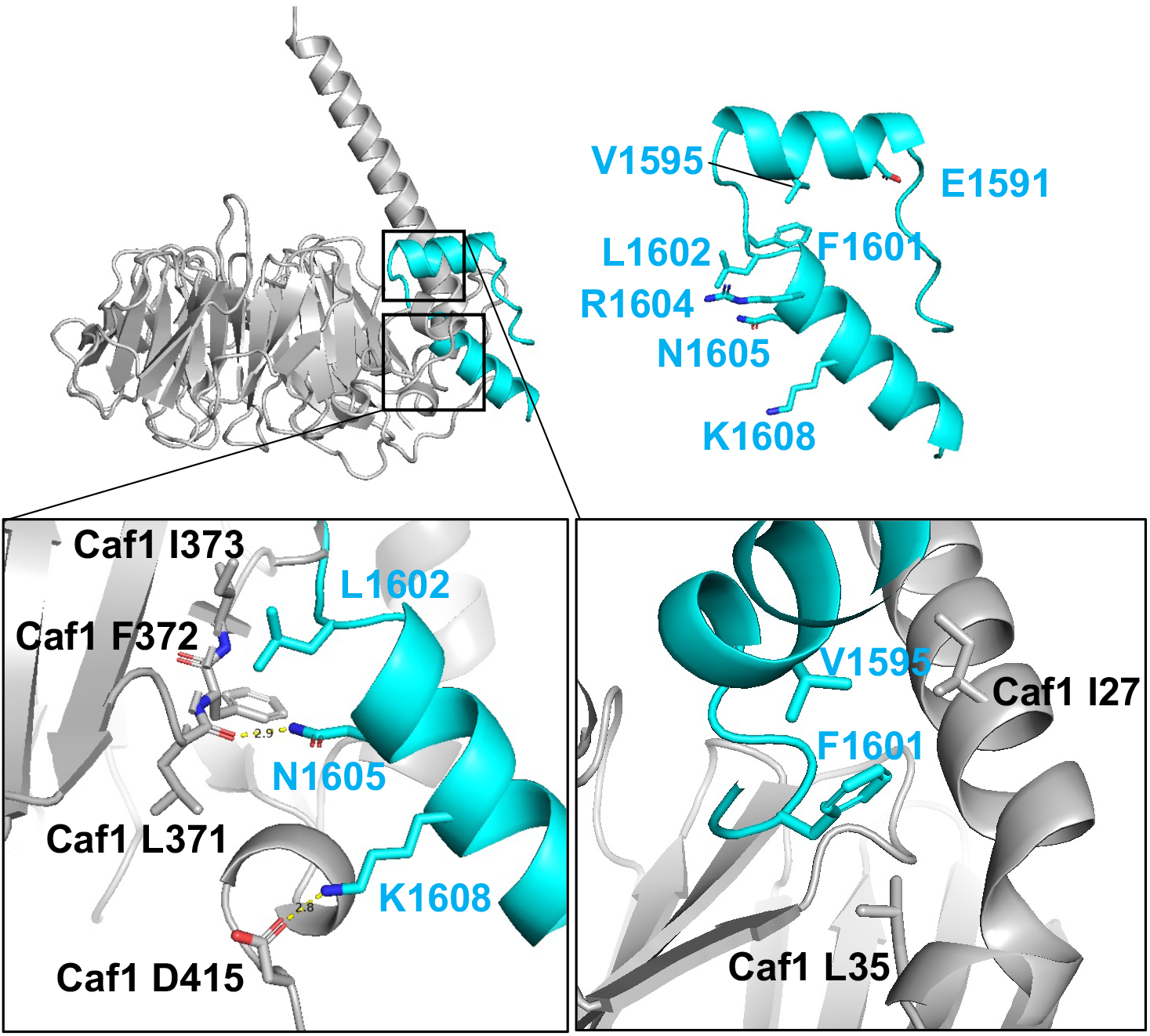
Conserved residues on Ash1_CBM extensively coordinate with Caf1 H4 binding pocket. Molecular interaction between L1602, N1605, and K1608 on Ash1. L1602 is found in close proximity to hydrophobic residues on Caf1 I373 and F372. N1605 and K1608 create salt bridges with the Caf1 L371 backbone carboxyl group, and the D415 side chain carbonyl group, respectively. In addition, V1595 and F1601 on Ash1 interact with Caf1 residues. V1591, F1601 and Caf1 I27, I35 side chains positioned in close proximity to stablize the Ash1-Caf1 binding via hydrophobic interaction.

**Figure S5.**
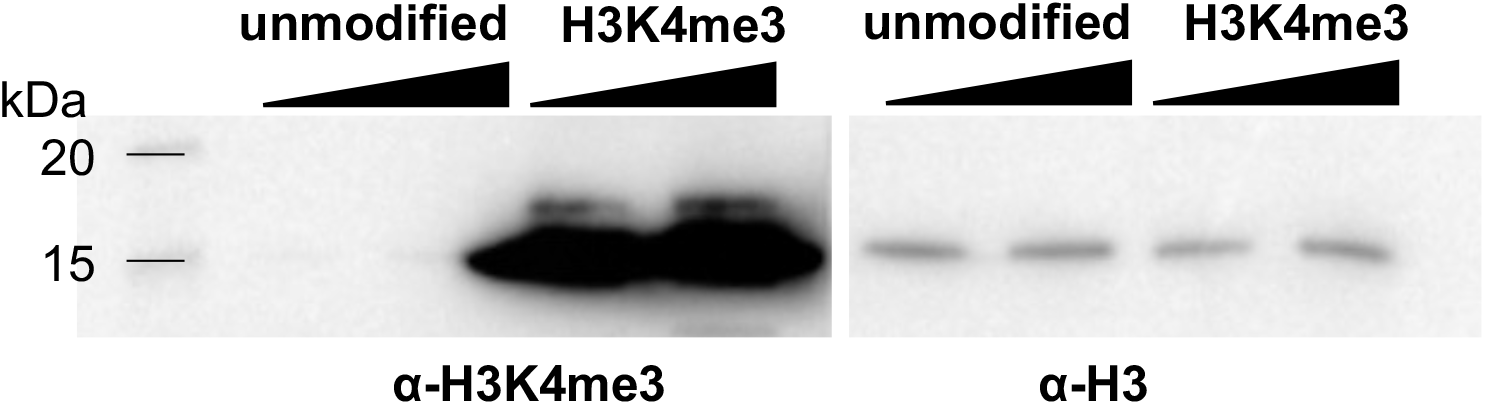
Western blot of MLA crosslinked H3K4me mimic nucleosome. Western blot data of unmodified and H3K4me3 MLA nucleosomes using α-H3K4me3 antibody and α-H3 antibody.

**Figure S6.**
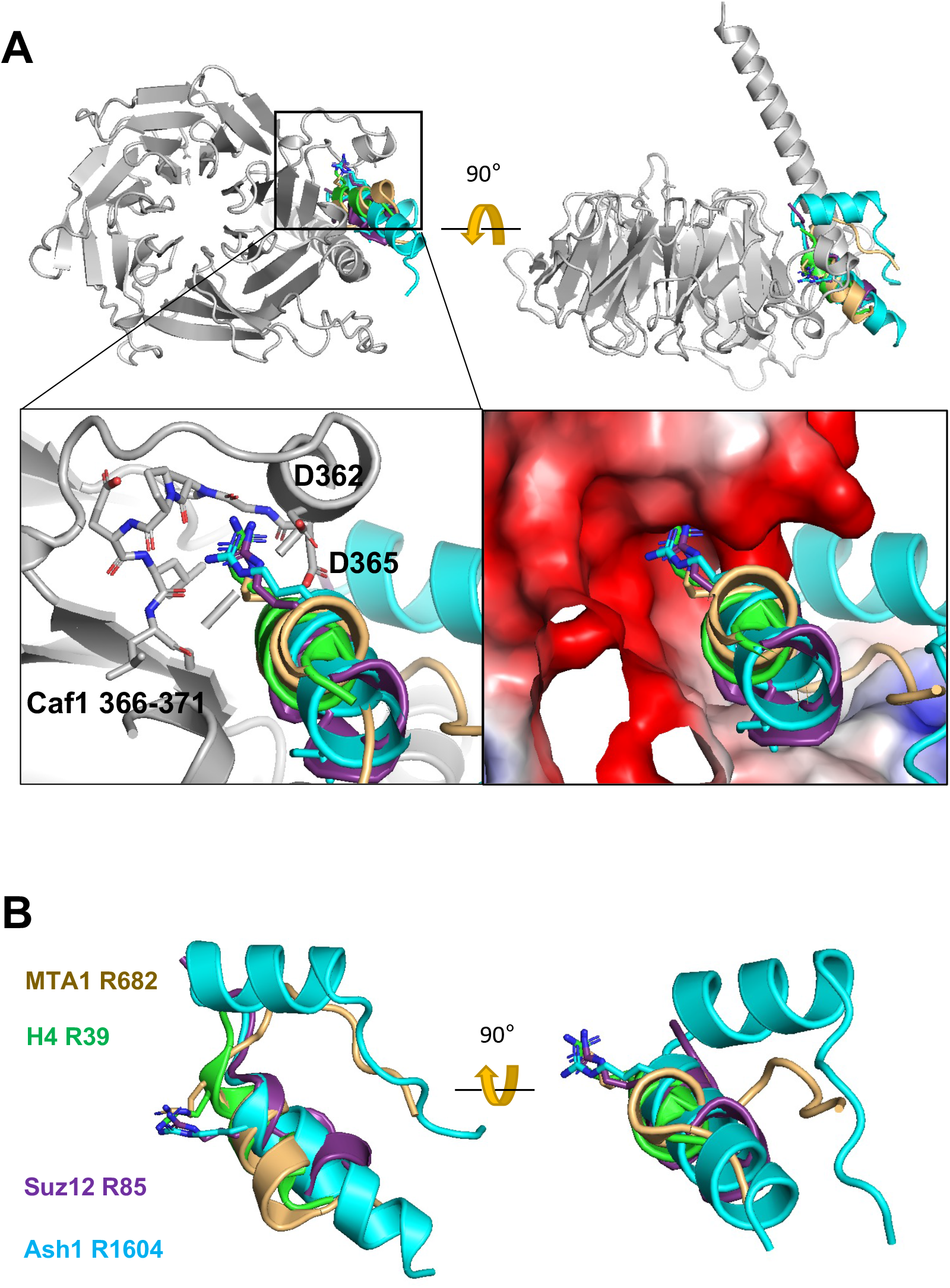
Proteins, which bind to the Caf1 H4 pocket, has conserved Arg. **A**. Superimposition of crystal structures of Caf1 perimeter binding proteins : H4 (green, PDB ID : 3c9c), MTA1 (yellow, PDB ID : 4pc0), Suz12 (purple, PDB ID : 2yb8) and the alphafold modelled CBM (cyan). D362 and D365 residues in Caf1 and the loop (366-371 a.a.) coordinate with the conserved Arg. **B**. The orientation of conserved Args in the Caf1 binding proteins.

